# Preparation of functional metagenomic libraries from low biomass samples using METa assembly and their application to capture antibiotic resistance genes

**DOI:** 10.1101/2024.06.29.601325

**Authors:** HM Allman, EP Bernate, E Franck, FJ Oliaro, EM Hartmann, TS Crofts

## Abstract

A significant challenge in the field of microbiology is the functional annotation of sequence novel genes from microbiomes. The increasing pace of sequencing technology development has made solving this challenge in a high-throughput manner even more important. Functional metagenomics offer a sequence-naïve and cultivation-independent solution. Unfortunately, most methods for constructing functional metagenomic libraries require large input masses of metagenomic DNA, putting many sample types out of reach. Here, we show that our functional metagenomic library preparation method, METa assembly, can be used to prepare useful libraries from much lower input DNA quantities. Standard methods of functional metagenomic library preparation generally call for 5 μg to 60 μg of input metagenomic DNA. We demonstrate that the threshold for input DNA mass can be lowered at least to 30.5 ng, a three-log decrease from prior art. We prepared functional metagenomic libraries using between 30.5 ng and 100 ng of metagenomic DNA, and found that, despite their limited input mass, they were sufficient to link MFS transporters lacking substrate-specific annotations to tetracycline resistance and to capture a gene encoding a sequence novel GNAT family acetyltransferase that represents a new streptothricin acetyltransferase, *satB*. Our preparation of functional metagenomic libraries from aquatic samples and a human stool swab demonstrate that METa assembly can be used to prepare functional metagenomic libraries from microbiomes that were previously incompatible with this approach.

## INTRODUCTION

Microbiomes, assemblages of bacteria, fungi, viruses and other microorganisms, are host to some of the highest levels of genetic diversity on the planet. Due to the difficulty in culturing all but a select low percentage of organisms in microbiomes (often estimated to be ∼0.1% to 1%) current methods for studying microbiomes rely on DNA extracted from these communities, metagenomic DNA^1–4^. High-throughput sequencing of metagenomic DNA provides unrivaled insights into the taxonomic structure of a microbial community and can shed significant light on its functional capacity as well^3^. However, even with the sequencing revolution, the genetic diversity of microbiomes is still beyond our ability to accurately annotate novel microbial genes. Complementing homology and structure-based methods for assigning function to novel genes is the high-throughput, sequence- and cultivation-naïve method of functional metagenomics. In this method, functional metagenomic libraries, sometimes referred to as shotgun cloning libraries, are prepared by fragmenting metagenomic DNA and cloning those fragments *en masse* into an expression microorganism. Once a library is prepared, any genes encoded within captured metagenomic DNA fragments are in theory able to be expressed by the host cell to give a phenotype that can be selected or screened for^4–7^ (**Figure 1A**). Functional metagenomic libraries can be divided into two categories based on the average size of their captured metagenomic DNA fragments, with small insert libraries generally capturing fragments between 2 kb and 10 kb in length and large insert libraries (*i.e,* cosmid, fosmid, and bacterial artificial chromosomes [BAC]) capturing fragments between 15 kb and 100s of kb in length^7^.

**Figure 1.**
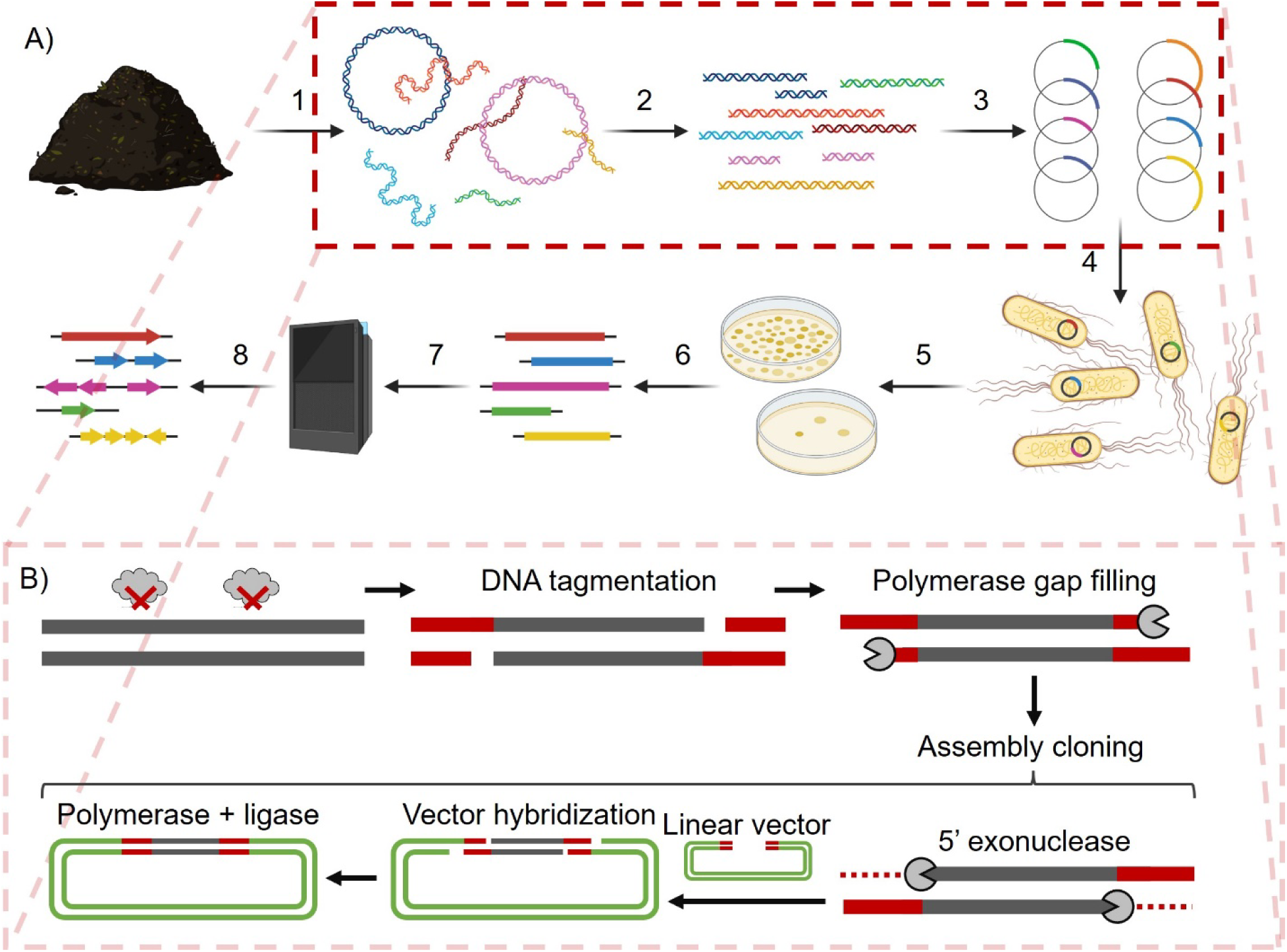
Functional metagenomics and METa assembly. A) The general steps in the preparation and application of a functional metagenomic library consist of: 1) Extraction of metagenomic DNA from a source microbiome. 2) Fragmentation of metagenomic DNA to preferred size range. 3) Packaging of inserts into expression vectors. 4) Transformation of host cells with vector library. 5) Screen or selection of functional metagenomic library for a phenotype of interest. 6) Collection of screened or selected metagenomic fragments. 7) Sequencing of selected inserts. 8) Open reading frame calling and annotation to identify potential genes underlying phenotypes of interest. B) Steps 2) and 3) above are modified in Mosaic ends tagmentation (METa) assembly. Fragmentation is achieved using Tn5 transposase tagmentation with mosaic end sequence oligos. Tagmented DNA is gap filled by polymerase and directly cloned (without amplification) into an expression vector with matching mosaic end sequences defining the cloning site using assembly cloning.

While functional metagenomics is a powerful method for linking novel genes to functions without requiring the growth of non-model organisms in the lab, it has its own shortcomings. These issues include the potential for gene toxicity, limited expression of foreign genes in library hosts such as *E. coli*, limited coverage of target metagenomes, and requirements for large quantities of metagenomic DNA for preparation of the functional metagenomic library itself. We recently reported the development of a new method for preparing small insert functional metagenomic libraries that we call METa assembly^8^. This functional metagenomic library preparation method relies on tagmentation to fragment input metagenomic DNA and an assembly-based cloning step to replace blunt ligation (**Figure 1B**). We previously demonstrated that this method circumvents one shortcoming of functional metagenomics, limited metagenomic coverage, by showing that METa assembly libraries are more efficient at capturing DNA and can produce approximately 80-fold larger libraries per input DNA mass. We also showed that METa assembly could be used to prepare functional metagenomic libraries encoding tens of gigabase pairs (Gb) of captured DNA using sub-μg inputs of metagenomic DNA, far lower than the standard 5 μg to 20 μg input^8,9^.

Here we build on our earlier results by testing if METa assembly can be used to prepare functional metagenomic libraries with even lower quantities of input DNA. We found that we could consistently prepare functional metagenomic libraries starting with 50 ng or less input DNA or even DNA extracted from a swab dipped into a 10 mg/ml human stool slurry. These libraries were large enough that, when selected for resistance to the antibiotics tetracycline and nourseothricin, they yielded genes predicted to encode novel proteins. These findings suggest that even low biomass microbiomes and samples can be studied by functional metagenomics using METa assembly.

## METHODS

### Strains and materials

Routine cultivation of *E. coli* was performed at 37°C with shaking at 250 rpm for aeration in the case of liquid cultures. Cultivation for cloning and starter cultures was performed using lysogeny broth (Miller) (LB) with the addition of kanamycin at 50 μg/ml to maintain plasmids. Antimicrobial susceptibility testing was performed using Cation-Adjusted Mueller-Hinton media (MH) (Teknova, 101320-364). *E. coli* clones used in this study are of the DH10B lineage and carry plasmids derived from pZE21 unless otherwise noted^8,10,11^. *E. coli* clones were maintained at -80°C in 15% glycerol in LB. Transposase enzymes and mosaic ends oligos were previously prepared and stored at -20°C^8,12^. Standard chemicals used in this study were of high purity for molecular biology and microbiology. Kanamycin sulfate (VWR Life Science, 75856-68), nourseothricin (Research Products International, N51200-1.0), tetracycline (Aldon Corp, TT0070-5GR), chloramphenicol (Thomas Scientific, 216035-25G), ciprofloxacin hydrochloride (Corning, 61-277-RG), azithromycin dihydrate (TCI, A2076), and florfenicol (Fisher Scientific, F08115G) were stored at 4°C as powders. Working solutions were stored at -20°C as filtered aqueous solutions at 50 mg/ml concentration (kanamycin and nourseothricin), 5 mg/ml in ethanol (tetracycline), 50 mg/ml in ethanol (chloramphenicol), 10 mg/ml in alkaline water (ciprofloxacin), or 50 mg/ml in DMSO (azithromycin and florfenicol).

### Preparation of 50 ng test functional metagenomic libraries

To determine if agarose extraction could be used to purify and size-select DNA following low mass tagmentation reactions, we used a DNA ladder (ThermoFisher Scientific 1 kb plus, 10787018) to mimic fragmented DNA. Mock tagmentation reactions were made with a final DNA concentration of 50 ng/μl by combining 1 μl of diluted DNA ladder with 3 μl of autoclaved milliQ water and 1 μl of 5X Taps tagmentation buffer (50 mM TAPS pH 8.5, 25 mM MgCl_2_, 50% v/v DMF in water)^12^. We added 0.25 μl of 1% sodium dodecyl sulfate (SDS) to the 5 μl mock reactions to simulate reaction quenching followed by 1 μl of 6x DNA dye (15% Ficoll 400 w/v, 0.25% Orange G w/v in water) before loading on to a 0.7% agarose gel in TAE buffer containing SybrSafe dye (ThermoFisher Scientific, S33102). DNA bands were visualized over blue light and a clean razor was used to cut out gel portions containing bands greater than 2 kb in length. DNA was extracted from the agarose gel fragments using a DNA extraction kit (New England Biolabs, T1020L) following manufacturer instructions with modification. Each agarose fragment was weighed and incubated with four volumes of gel dissolving buffer for at least 10 minutes at 55°C. The dissolved agarose was then applied to a spin column, spun through, then re-applied a second time. The standard protocol was followed until the elution step where each column was eluted twice sequentially with 6 μl of 55°C autoclaved milliQ water. DNA concentration was measured using the Quant-It DNA quantification system (Invitrogen).

To test the preparation of functional metagenomic libraries using 50 ng of input metagenomic DNA, we used a sample of previously extracted metagenomic DNA from a Canada goose fecal pellet that was stored at -20°C^8^. Briefly, metagenomic DNA for that sample was extracted using a DNeasy PowerSoil kit (Qiagen, catalog no. 12888-100) with modifications including incubation of fecal material in Qiagen buffer CD1 at 65°C for 10 minutes and 95°C for 10 minutes^8,13^. For library preparation, we followed our previously published METa assembly protocol^8^. Briefly, triplicate 5 μl tagmentation reactions containing 50 ng of metagenomic DNA, 1x TAPS-DMF buffer, and 25 ng of in-house transposase^12^ loaded with mosaic end sequence oligos were incubated at 55°C for 7 minutes followed by quenching by addition of SDS to reach 0.05% final concentration and incubation for 5 more minutes at 55°C. To each reaction, 2 μl of 6x loading dye was added and fragmented metagenomic DNA in the range of 2 kb to 9 kb was extracted by gel excision as described above. The 12 μl of size-selected metagenomic DNA fragments were gap-filled by the addition of 12 μl of 2x Q5 DNA polymerase (New England Biolabs, M0492S) (pre-incubated at 98°C for 30 seconds) and held at 72°C for 15 minutes before purification by PCR and DNA cleanup kit (New England Biolabs, T1030S) according to manufacturer instructions with the following modifications: DNA dissolved in two volumes of binding buffer was applied to the spin column and the flow-through was applied a second time. At elution, we used two sequential applications of 6 μl of 55°C autoclaved milliQ water.

Metagenomic DNA inserts were cloned into our plasmid pZE21-ME using 2x NEBuilder HiFi assembly mix (New England Biolabs, E2621L) according to manufacturer instructions (assuming an insert DNA mass of 15 ng and an average fragment size of 2 kb). The triplicate libraries were purified by PCR and DNA cleanup kit and the full 12 μl of purified library DNA was introduced into 25 μl of *E. coli* DH10B electrocompetent cells (New England Biolabs, C3020K) by electroporation at 1.8 kV in 0.1 mm cuvettes. Actual voltage reached and the electroporation time constant were recorded for all electroporation reactions to detect failed or low efficiency transformations. All three libraries were rescued post-transformation by inoculation into 1 ml of SOC outgrowth medium (New England Biolabs, B9020S) pre-warmed to 37°C and incubated with aeration at 37°C for 1 hour.

Following recovery, 100 μl of 100-fold, 10,000-fold, and 1,000,000-fold diluted cultures were plated on to LB agar with kanamycin and incubated overnight at 37°C. The colonies were counted to estimate the number of unique clones resulting from each transformation. The remaining cells (∼1 ml) were inoculated into 50 ml of LB with kanamycin and incubated at room temperature overnight with aeration to amplify the functional metagenomic library. Once cultures reached a 1 cm pathlength optical density at 600 nm (OD_600_) of between 0.6 and 1.0, cells were collected by centrifugation at 4,000 rcf for 7 minutes at 4°C. Pellets were resuspended in 10 ml of LB containing kanamycin and 15% glycerol and stored at -80°C as 1 ml aliquots. The triplicate functional metagenomic library sizes were determined using the Dantas lab calculator (http://dantaslab.wustl.edu/LibSizeCalc/) considering unique cell count, determined above, average metagenomic insert size, and the proportion of clones containing an empty plasmid. This proportion and the average insert size were determined by performing colony PCR as previously described on 9 clones from each replicate library^8^.

### Preparation of an aquatic functional metagenomic library from a Shedd Aquarium microbiome

Preparation of a functional metagenomic library from aquatic metagenomic DNA began with a previously extracted metagenomic DNA sample. The initial sample was collected from the Shedd Aquarium in Chicago in the Underwater Beauty exhibit in May, 2019 and metagenomic DNA was extracted^14^. A functional metagenomic library was prepared from 50 ng of DNA as described above and evaluated by colony counting and colony PCR (n=14 colonies) to determine library size. After preparation of the functional metagenomic library, quantification of the input DNA by fluorescent dye (Quant-It) revealed that only 30.5 ng of metagenomic DNA had been used as input for this library and this corrected DNA mass was used in determining library preparation efficiency.

### Functional metagenomic selection for antibiotic resistance from the aquarium functional metagenomic library

The aquarium functional metagenomic library was selected for resistance to the antibiotics tetracycline, ciprofloxacin, and florfenicol. MH agar plates containing 50 μg/ml kanamycin were prepared to contain each antibiotic at the following concentrations previously established to be inhibitory to *E. coli*^15^: 8 μg/ml tetracycline, 0.5 μg/ml ciprofloxacin, and 8 μg/ml florfenicol. An aliquot of the functional metagenomic library was thawed from -80°C storage and a volume of suspension calculated to contain approximately 10-fold more total cells than the unique cell count was plated on each antibiotic and incubated overnight at 37°C. A parallel series of agar plates were plated with a similar cell count of *E. coli* cells containing only empty vectors as a control.

Following incubation, 15 colonies each from the tetracycline and florfenicol selections were collected and re-streaked onto LB agar plates containing kanamycin and either tetracycline or florfenicol to obtain pure cultures of resistant clones. Plasmids were extracted from each clone by miniprep kit (New England Biolabs, T1010L) and whole plasmid sequencing was performed by Plasmidsaurus using Oxford Nanopore Technology with custom analysis and annotation. The metagenomic inserts were identified by the presence of flanking mosaic end sequences (5’-agatgtgtataagagacag-3’) and clustered on the Clustal Omega webserver^16^ to de-replicate metagenomic fragments that were captured in the selection more than once. The metagenomic inserts targeted for further study (TET1, TET3, TET13) were analyzed by the MetaGeneMark program^17^ to identify potential open reading frames which were subjected to analysis by Basic Local Alignment Search Tool (BLAST)^18,19^ against the NCBI non-redundant database and The Comprehensive Antibiotic Resistance Database (CARD)^20,21^. The annotated metagenomic DNA fragments were made into schematics using ApE plasmid editor^22^ and Microsoft PowerPoint.

### Preparation of a functional metagenomic library from a fecal swab

An aliquot of the ZymoBIOMICS fecal reference standard (Zymo Research, D6323) was used as a source for a fecal microbiome swab functional metagenomic library. An aliquot of fecal standard was thawed from -80°C storage and a sterile swab was dipped into the tube and gently wrung out against the tube’s side as it was removed. Metagenomic DNA was extracted from the swab using a PowerSoil Pro kit (Qiagen, 47014). The swab was agitated in 800 μl of C1 solution to release material and DNA extraction was performed following manufacturer instructions with the eluted DNA reduced in volume using a vacuum concentrator. A functional metagenomic library was prepared by METa assembly as above, with the tagmentation reaction scaled to match the 100.8 ng of metagenomic DNA available from the extraction. Library size was evaluated by colony count and colony PCR as above (n=14 colonies).

### Functional metagenomic selection for nourseothricin resistance from the stool functional metagenomic library

In our previous nourseothricin resistance selection^8^, we used agar plates containing 64 μg/ml nourseothricin but this concentration of drug is likely well beyond the *E. coli* minimal inhibitory concentration (MIC). To determine conditions appropriate for a functional metagenomic selection for nourseothricin resistance genes, we performed an agar dilution assay on MH agar with drug concentrations ranging from 0.5 μg/ml to 2048 μg/ml in two-fold jumps^23^. Each agar plate was inoculated with 100 μl of a 0.5 McFarland unit suspension of *E. coli* DH10B harboring an empty pZE21-ME plasmid. Plates were incubated at 37°C for roughly 24 hours prior to being observed for growth to determine an appropriate concentration for functional metagenomic selections.

A functional metagenomic selection for nourseothricin resistance genes was carried out as above with the stool swab functional metagenomic library plated to carry an estimated ten-fold excess of cells over unique clones. The functional metagenomic library was plated on MH agar with 16 μg/ml nourseothricin and 50 μg/ml kanamycin. In parallel, a suspension of a control *E. coli* DH10B clone with empty vector was also plated as above at a similar predicted titer. The plates were incubated at 37°C overnight. A single colony on the nourseothricin plate was picked and re-streaked to obtain a pure culture and processed by miniprep kit to obtain a plasmid for sequencing. The plasmid sequence was analyzed as before. The predicted acetyltransferase open reading was synthesized into a pZE21-derivative plasmid^10,11^ by Integrated DNA Technologies (IDT) and an aliquot of electrocompetent *E. coli* DH10B was transformed with the plasmid for further resistance study.

### Phylogenetic analyses

For phylogenetic analysis of the putative tetracycline efflux pumps (TET1, TET3, and TET13) we downloaded CARD amino acid sequences^20,21^ for the ontology term “major facilitator superfamily (MFS) antibiotic efflux pump” and added the functionally-selected predicted efflux pump amino acid sequences to the FASTA file. For analysis of the predicted nourseothricin resistance protein, we downloaded CARD protein sequences for the ontology term “acylation of antibiotic conferring resistance”, which includes the streptothricin, aminoglycoside, apramycin, virginiamycin, capreomycin, and chloramphenicol acetyltransferase sequences. We also downloaded the amino acid sequence for the NAT nourseothricin acetyltransferase^24^.

Phylogenetic trees were prepared by first aligning the protein sequences using the Clustal Omega web server with default settings^16^. Next a maximum likelihood phylogenetic tree was prepared from the alignment using IQ-TREE^25,26^ with the “Find best and apply” substitution model, standard 100 replicates bootstrap and otherwise default settings. The resulting consensus trees were visualized using the Interactive Tree of Life (iTOL) display and annotation tool^27^.

The streptothricin acetyltransferase multiple protein alignment was prepared using Clustal Omega as above using only the STAT, NAT, SAT-2, SAT-3, SAT-4, SatA, and SatB amino acid sequences. This also provided a percent identity matrix for the streptothricin acetyltransferase proteins. The alignment was edited and visualized in Jalview^28^ and amino acid residue numbering was set according to the *Bacillus anthracis* SatA crystal structure sequence^29^.

### Microbroth dilution antimicrobial susceptibility testing

The antimicrobial resistance of *E. coli* clones expressing metagenomic DNA fragments (TET1, TET3, TET13, and NTC1), the *satB* open reading frame, or the *stat* gene (pGDP1 stat was a gift from Gerard Wright (Addgene plasmid # 112886; http://n2t.net/addgene:112886; RRID:Addgene_112886)) were quantified by microbroth dilution assay^23,30^. Briefly, the corresponding clones and control *E. coli* carrying an empty plasmid were streaked for single colonies on LB agar with kanamycin. Approximately 10 colonies for each clone were suspended in 2 ml of MH media with kanamycin and brought to an OD_600_ of between 0.1 and 0.15 to correspond to a 0.5 McFarland standard density, then diluted 100-fold in MH broth with kanamycin. We prepared a 96-well plate (Costar, 3370) to contain 50 μl of MH broth with 50 μg/ml kanamycin and 2-fold increasing concentrations of tetracycline, chloramphenicol, azithromycin, or nourseothricin at twice the final target concentration. We added 50 μl of the diluted bacterial suspensions to this plate, with each clone assayed in quadruplicate. The 96-well plates were sealed with a Breathe-Easy membrane (MilliporeSigma, Z380059) and incubated at 37°C with shaking for 20 to 24 hours. Culture density was recorded as OD_600_ with a 1 cm pathlength correction using an Epoch 2 plate reader (BioTek) and concentration-response curves were fitted to four parameter Hill equations in GraphPad Prism 10.2.3 (GraphPad Software, La Jolla, CA, USA) to determine 50% inhibitory concentration (IC_50_) values. Differences between IC_50_ values for control *E. coli* and the tested clones were evaluated for significance in GraphPad Prism using lognormal Brown-Forsythe ANOVA tests with multiple comparison correction using the Dunnett T3 method.

### SatB structural prediction

Structural modelling of the SatB protein was performed using AlphaFold2 through the ColabFold v1.5.5 Jupyter notebook^31,32^. The predicted amino acid sequence was entered as query sequence and default parameters were used to generate the model. The model was superimposed against the SatA crystal structure (3PP9)^29,33,34^ using the Mol* 3D viewer^35^ hosted by the Research Collaboratory for Structural Bioinformatics Protein Data Bank (RCSB PDB)^36^. Mol* 3D calculated the root mean squared deviation (RMSD) of the resulting superposition of SatB on SatA.

### Sequence availability

Sequences for the TET1, TET3, TET13, and NTC1 metagenomic DNA fragments and the proposed *satB* gene can be found under GenBank accession numbers PV822507, PV822508, PV822509, PV8225110, and PV822511.

## RESULTS

### METa assembly can reproducibly prepare Gb sized libraries with 50 ng of input metagenomic DNA

Our published METa assembly protocol tagments metagenomic DNA at a concentration of 10 ng/μl, meaning a 50 ng DNA tagmentation reaction would have a 5 μl volume where accurate pipetting starts to become difficult. To test the practicality of extracting useful DNA fragments from this size reaction, we diluted 50 ng of a DNA ladder in tagmentation buffer in triplicate to a concentration of 10 ng/μl to make mock 5 μl tagmentation reactions. The triplicate mock fragmented samples were loaded on to an agarose gel for excision of bands ranging from approximately 2 kb to 10 kb (**Supplemental figure 1A**). This resulted in recovered DNA quantities (acting as stand-ins for fragmented metagenomic DNA) averaging 12.7 ng (± 2.7 standard deviation). Given that the total mass of the DNA bands 2 kb and higher was 16 ng, this represented a 79% yield and suggested that size-selection of fragmented DNA from a 50 ng tagmentation reaction could be feasible (**Supplemental figure 1B**).

Next, we tested this explicitly by preparing a 50 ng metagenomic DNA functional metagenomic library in triplicate. Using previously extracted goose fecal metagenomic DNA as input^8^, we performed three tagmentation reactions in parallel and used gel excision to size-select for DNA fragments between 2 kb and 10 kb (**Supplemental figure 2A**). These fragments were used to prepare three functional metagenomic libraries by METa assembly. During the electroporation step, replicates 1 and 3 reached full voltage (1.8 kV) and had time constants of 4.6 and 4.8 seconds respectively. Replicate 2 reported a voltage of 1.65 kV and a 1.0 second time constant, consistent with the sample ‘sparking’. Replicate #1 had an average insert size of 2.48 kb, no empty vectors, and an average library size of 0.89 Gb with a range of 0.69 Gb to 1.1 Gb. For replicate #2 these values were 2.62 kb, no empty vectors, and 0.1 Gb with a range of 0.08 Gb to 0.13 Gb. Replicate #3 values were 3.13 kb, no empty vectors, and 2.46 Gb with a range of 1.61 Gb to 3.27 Gb. Averaging across the three libraries we found an overall average insert size of 2.75 kb with no empty vectors (**Supplemental figure 2B**), an average library size of 1.15 Gb, and a library preparation efficiency of 23 Gb/μg of input DNA. Notably, these averages include replicate #2 with a dramatically smaller library size (100 Mb compared to 890 Mb and 2.5 Gb) (**Table 1**). If replicate #2 is excluded, the average library size for the 50 ng functional metagenomic libraries increases to 1.68 Gb with an efficiency of 33.6 Gb/μg.

**Table 1.**
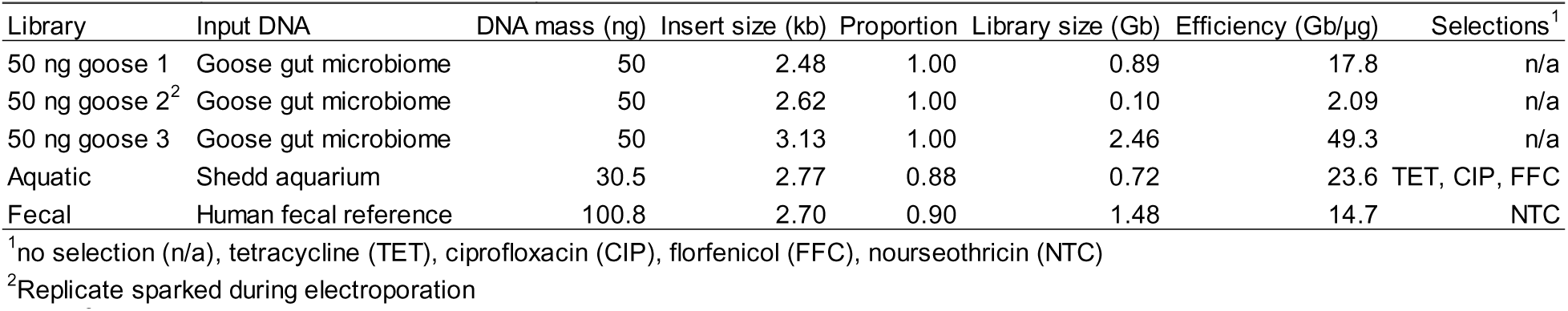
Summary statistics of functional metagenomic libraries.

### METa assembly preparation of a functional metagenomic library from a low biomass aquarium microbiome

Two areas where input mass is limiting for classic functional metagenomic library preparation are 1) when the microbiome of interest is low in biomass (*e.g.,* aquatic sources as opposed to soil or fecal samples) and 2) when the microbiome of interest is not readily available for re-sampling. We decided to test low-input mass METa assembly on a microbiome that fulfills both limitations. In a previous project, metagenomic DNA was extracted from the Underwater Beauty exhibit at the Shedd Aquarium in Chicago, Illinois following filtration capture of planktonic cells^14^. We used a sample of previously extracted metagenomic DNA to prepare a functional metagenomic library. Tagmentation was performed on 30.5 ng of metagenomic DNA and fragments from 2 kb to 10 kb were size selected by gel excision. Following transformation, random colonies from the functional metagenomic library were used as template for PCR to amplify captured metagenomic inserts to determine library statistics. We found that this aquatic functional metagenomic library captured 720 Mb of metagenomic DNA (range of 0.42 Gb to 1.17 Gb) with an average insert size of 2.77 kb and with 88% of plasmids containing an insert. The library preparation efficiency was calculated to be 23.6 Gb/μg (**Table 1**).

### Tetracycline resistance in an aquarium functional metagenomic library

To evaluate the usefulness of this functional metagenomic library, we used it to investigate antimicrobial resistance against three antibiotics that are commonly used in aquaculture: tetracycline, florfenicol, and ciprofloxacin^37,38^. The selection on ciprofloxacin did not yield any resistant colonies but colonies were recoverable from both the tetracycline and florfenicol plates. As an initial investigation, fifteen colonies from only the tetracycline selection were collected and submitted for whole plasmid sequencing. We dereplicated repeated inserts, which resulted from plating a ten-fold excess of total cells over unique cells, resulting in three *E. coli* clones with unique metagenomic DNA fragments we termed TET1, TET3, and TET13. The metagenomic DNA captured in these cones had top nucleotide BLAST hits to *Lentibacter algarum* strain SH36 (95% coverage, 82.09% identity), *Legionella pneumophila* strain NY23 (30% coverage, 66.87% identity), and *Saccharophagus degradans* strain FZY0027 (32% coverage, 70.26% identity), respectively.

We annotated the metagenomic DNA fragments captured in these three clones (**Supplemental figure 3**) and found them to each contain predicted MFS family efflux pumps with low full-length amino acid identity (<30%) to their closest match in The Comprehensive Antibiotic Resistance Database (CARD)^20,21^ (**Figure 2A**). When placed on a phylogenetic tree alongside known antibiotic resistance efflux pumps, TET1 and TET13 clustered with each other, a bicyclomycin efflux pump (BCR-1), and efflux pumps associated chloramphenicol resistance (PexA, MdfA, and Cml family pumps) (**Figure 2B**). TET3 instead clustered with efflux pumps with a variety of targets including tetracycline (TetV, TAP), macrolides (MefD), and multiple antimicrobials (Hp1181, EfmA,) (**Figure 2B**).

**Figure 2.**
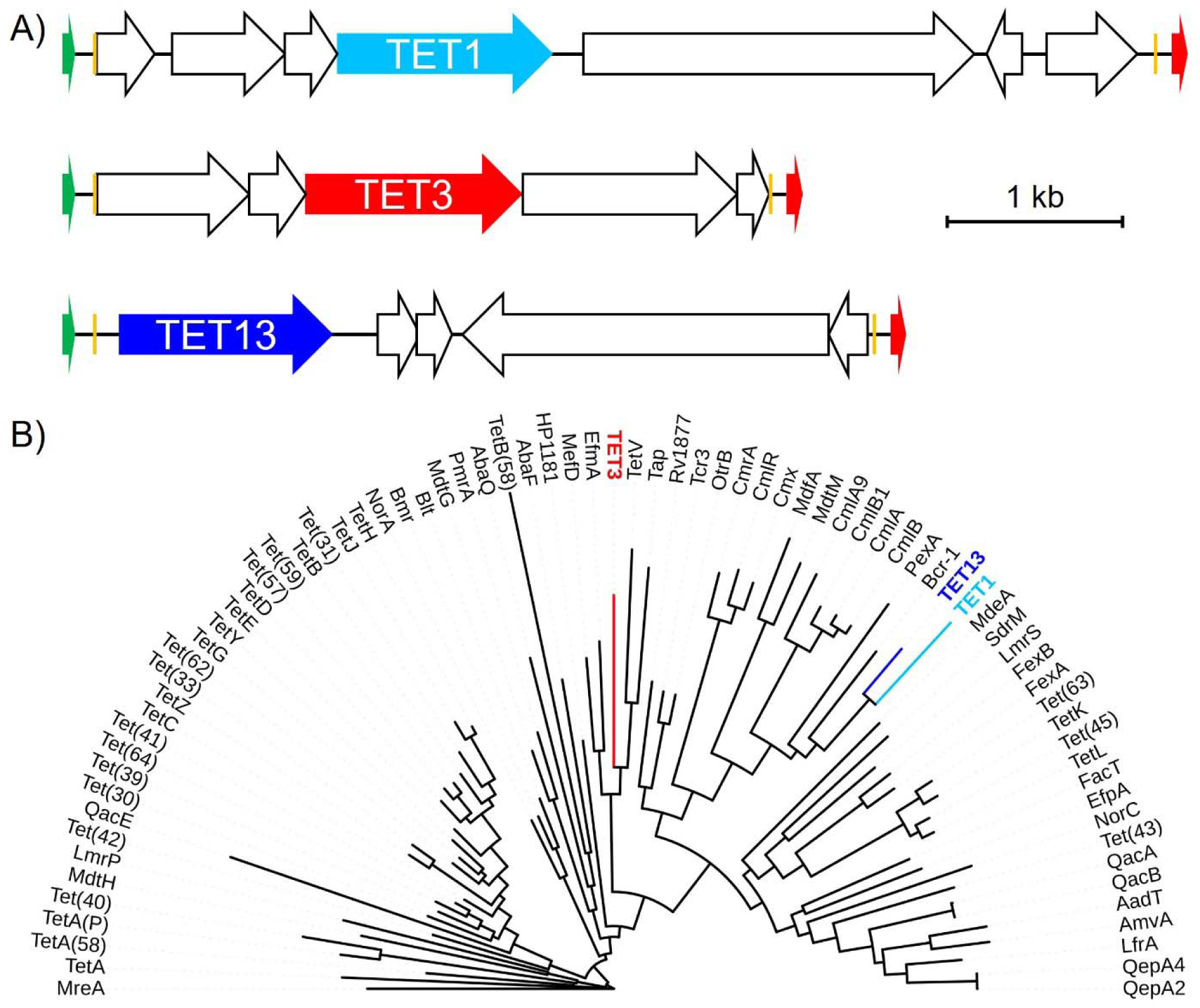
Tetracycline selected metagenomic DNA fragments from an aquarium microbiome. A) Gene schematics of the tetracycline-selected TET1, TET3, and TET13 metagenomic DNA fragments with predicted efflux pumps highlighted (TET1 teal, TET3 red, TET13 blue). Other predicted open reading frames are shown as empty arrows and plasmid backbone markers are highlighted as follows: Promoter (green), mosaic end sequences (orange), terminator (red). B) Maximum likelihood consensus phylogenetic tree of MFS efflux pumps with predicted efflux pump amino acid sequences from TET1 (teal), TET3 (blue), and TET13 (red) fragments highlighted.

The resistance of all three clones to tetracycline, chloramphenicol, and azithromycin were measured by microbroth dilution. All three clones showed increased resistance to tetracycline, with TET1 and TET3 notably showing an ability to grow in the presence of 8 μg/ml tetracycline, the breakpoint for resistance determination in *Enterobacterales* like *E. coli* (**Figure 3A**). All three *E. coli* clones gained a statistically significant increase in their tetracycline 50% inhibitory concentration (IC_50_) value (**Figure 3B**). Two of the clones showed a statistically significant change in chloramphenicol resistance (TET1 being more susceptible and TET13 being more resistant) (**Figure 3C and 3D**). However, the limited change in absolute terms (less that a two-fold change) suggests they are not biologically significant. None of the three clones showed a significant change in azithromycin resistance compared to the control clone (**Figure 3E and 3F**).

**Figure 3.**
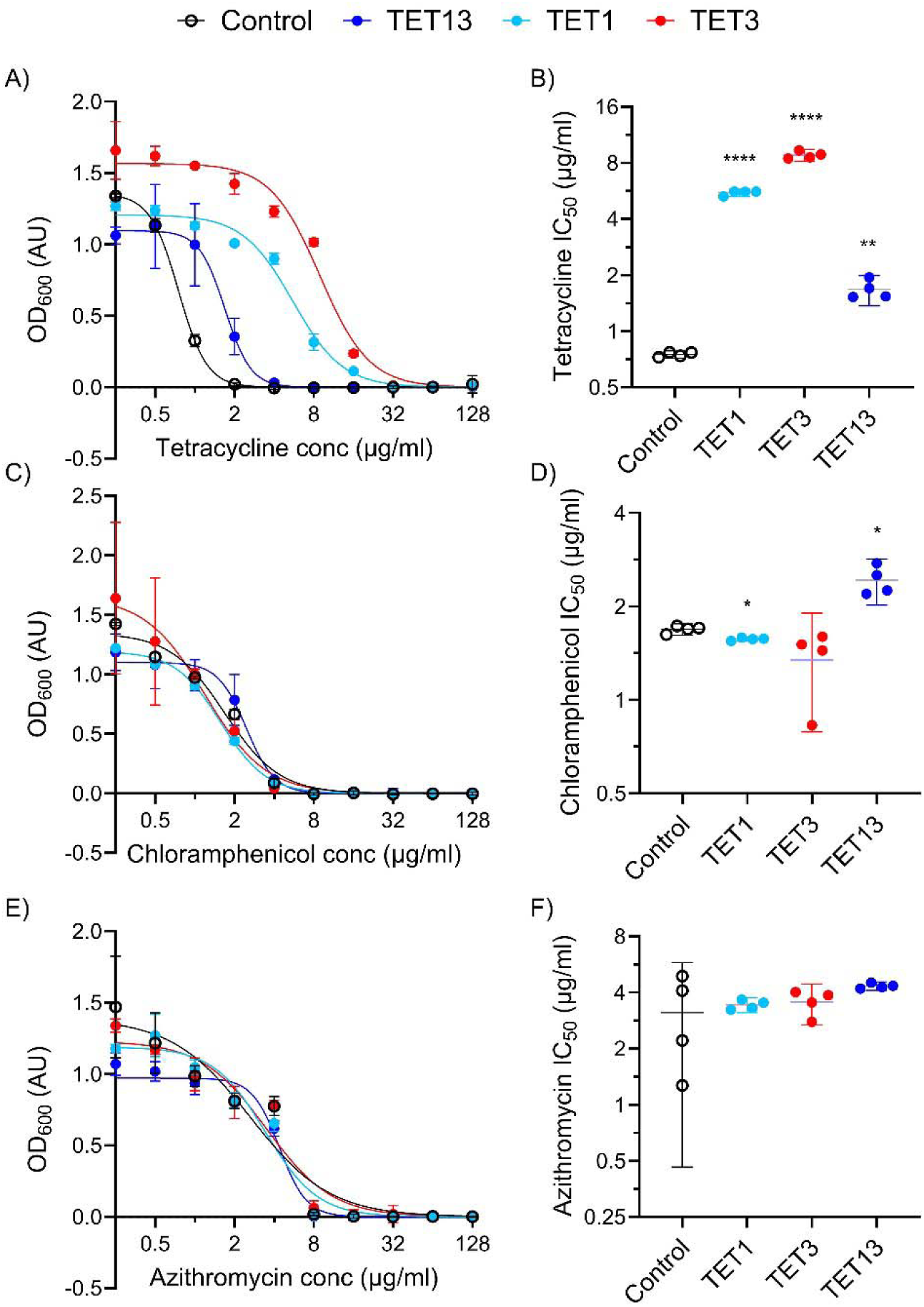
Efflux pump containing metagenomic DNA fragments from an aquarium confer tetracycline resistance. Microbroth dilution assay curves and calculated 50% inhibitory concentration (IC_50_) values for *E. coli* clones carrying the indicated metagenomic DNA fragments (TET1, TET3, or TET13). A) and B) tetracycline, C) and D) chloramphenicol, or E) and F) azithromycin. (**** p<0.0001, ** p<0.005, *p<0.05, n=4 for all).

### METa assembly preparation of a functional metagenomic library from a human stool swab

Another area where the ability to prepare a functional metagenomic library using minimal DNA resources could be beneficial is within clinical settings. For example, fecal swabs provide access to the medically important gut microbiome, but extraction of metagenomic DNA from this sample type falls far below what classic functional metagenomic library preparation methods call for^15,39^. To test the possibility of medical swabs being used as input for METa assembly, we turned to the ZymoBIOMICS fecal reference with TruMatrix technology fecal standard. Each aliquot of this standard is expected to contain 10 mg (wet weight) of human fecal material in 1 ml of DNA/RNA shield preservation buffer. We sampled this microbiome by dipping a sterile swab into a thawed tube of the material followed by DNA extraction from the swab, resulting in collection of 100.8 ng of DNA. We prepared a functional metagenomic library by METa assembly and evaluated it as above and found 90% of colonies contained an insert, each with an average size of 2.7 kb, for a total expected library size of 1.48 Gb (range of 0.62 Gb to 2.32 Gb) and a library preparation efficiency of 14.7 Gb/μg (**Table 1**).

### Nourseothricin resistance in a human stool functional metagenomic library

We performed a functional metagenomic selection for nourseothricin resistance in the human gut microbiome using the Zymo fecal library. In our previous work we performed nourseothricin selections using a concentration of 64 μg/ml^8^ but here we found that 16 μg/ml is more than sufficient. This selection resulted in the capture of a single nourseothricin resistant colony which we collected and extracted a plasmid from to sequence the resistance-conferring metagenomic DNA fragment. A parallel experiment in which an equal titer of control *E. coli* cells containing an empty plasmid was plated on agar containing 16 μg/ml nourseothricin resulted in the growth of no colonies.

Following whole plasmid sequencing of the resistant colony, we found that the NTC1 metagenomic DNA fragment showed limited identity and coverage with organisms in the NCBI database, with the top nucleotide BLAST hit being *Clostridia* bacterium i40-0019-1A8 (20% coverage, 76.08% identity). The Zymo fecal reference has been shotgun sequenced by other groups and we confirmed the presence of short reads in these sequencing runs (*i.e.,* SRX26261613^40^) with >98% nucleotide identity to the NTC1 metagenomic DNA fragment, confirming its origin in the Zymo stool material.

The metagenomic DNA fragment was predicted to contain three open reading frames including a potential GCN5-Related N-Acetyltransferase (GNAT) (**Figure 4A**), the enzyme family to which antibiotic-inactivating acetyltransferase enzymes belong. We prepared a phylogenetic tree with the predicted streptothricin acetyltranferase, which we termed SatB, alongside the six other streptothricin acetyltransferase family members (STAT, NAT, SatA, SAT-2, SAT-3, and SAT-4) and aminoglycoside, virginiamycin, capreomycin, and chloramphenicol acetyltransferases from the CARD database (**Figure 4B**). This tree places SatB in the same clade as the other streptothricin acetyltransferases but at a distance comparable to other distinct members of the family.

**Figure 4.**
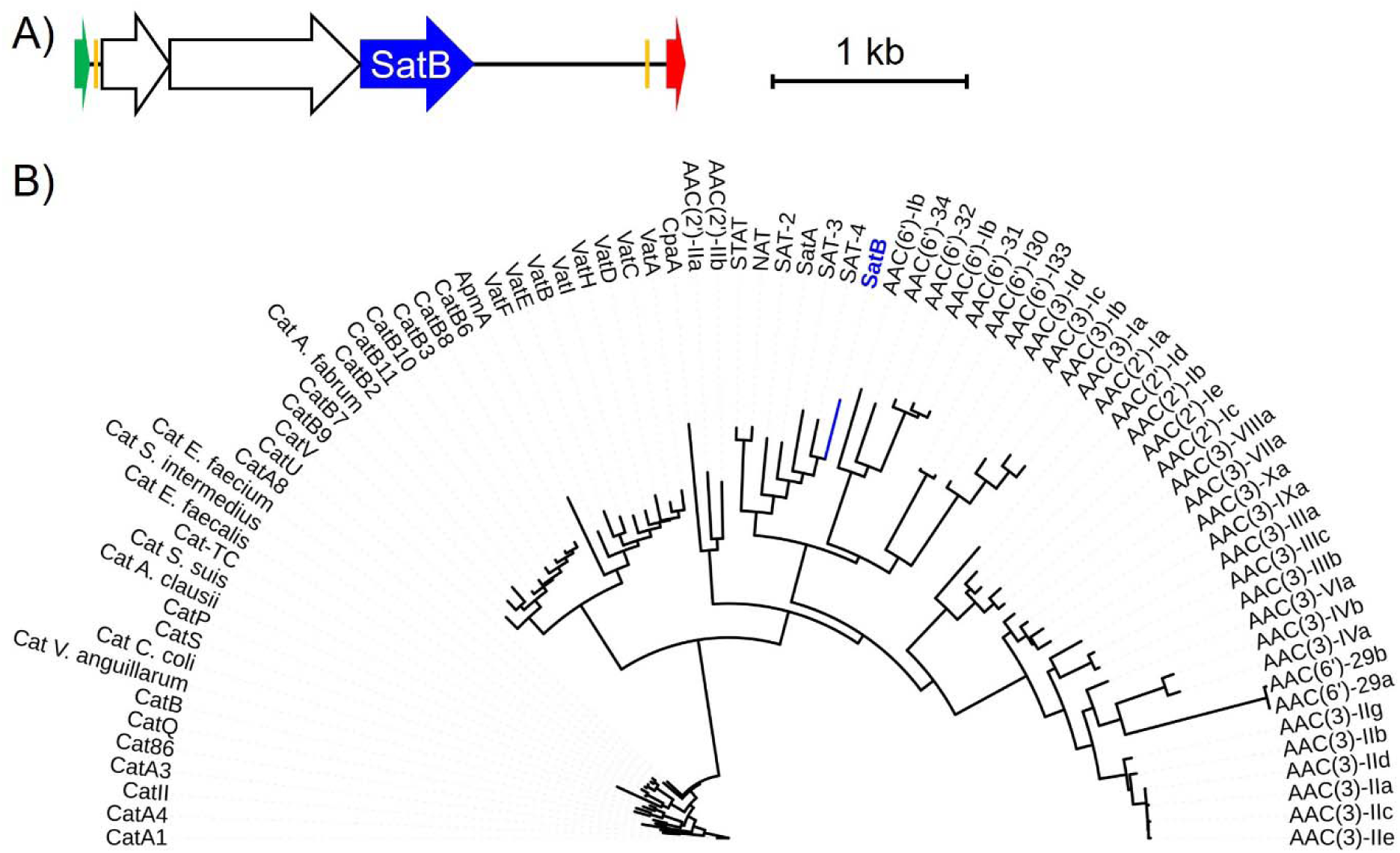
Predicted novel streptothricin acetyltransferase from the human gut microbiome. A) Gene schematics of the NTC1 metagenomic DNA fragment with the predicted acetyltransferase gene highlighted (‘*satB*’, blue). Predicted genes captured on the same DNA fragment are shown as empty arrows. Plasmid backbone elements are: Promoter (green), mosaic end sequence (orange), terminator (red). B) Maximum likelihood consensus phylogenetic tree of the predicted SatB protein (blue) in the context of known streptothricin acetyltransferases (SATs, NAT, and STAT) and aminoglycoside, chloramphenicol, virginiamycin, apramycin, and capreomycin acetyltransferases.

To explore this further we investigated the ability of the NTC1 metagenomic DNA fragment and of the predicted *satB* open reading frame to confer nourseothricin resistance in *E. coli*. As a positive control we also included an *E. coli* clone expressing the *stat* gene^41^. All three clones showed an ability to grow at higher nourseothricin concentrations than a control *E. coli* clone with an empty plasmid (**Figure 5A**). Expression of the NTC1 metagenomic DNA fragment was sufficient to increase the nourseothricin resistance of *E. coli* by approximately 64-fold and expression of the *stat* gene or the putative *satB* gene provided even greater resistance, allowing *E. coli* to grow at nourseothricin concentrations at least 1,000-fold greater than when carrying an empty plasmid (**Figure 5B**).

**Figure 5.**
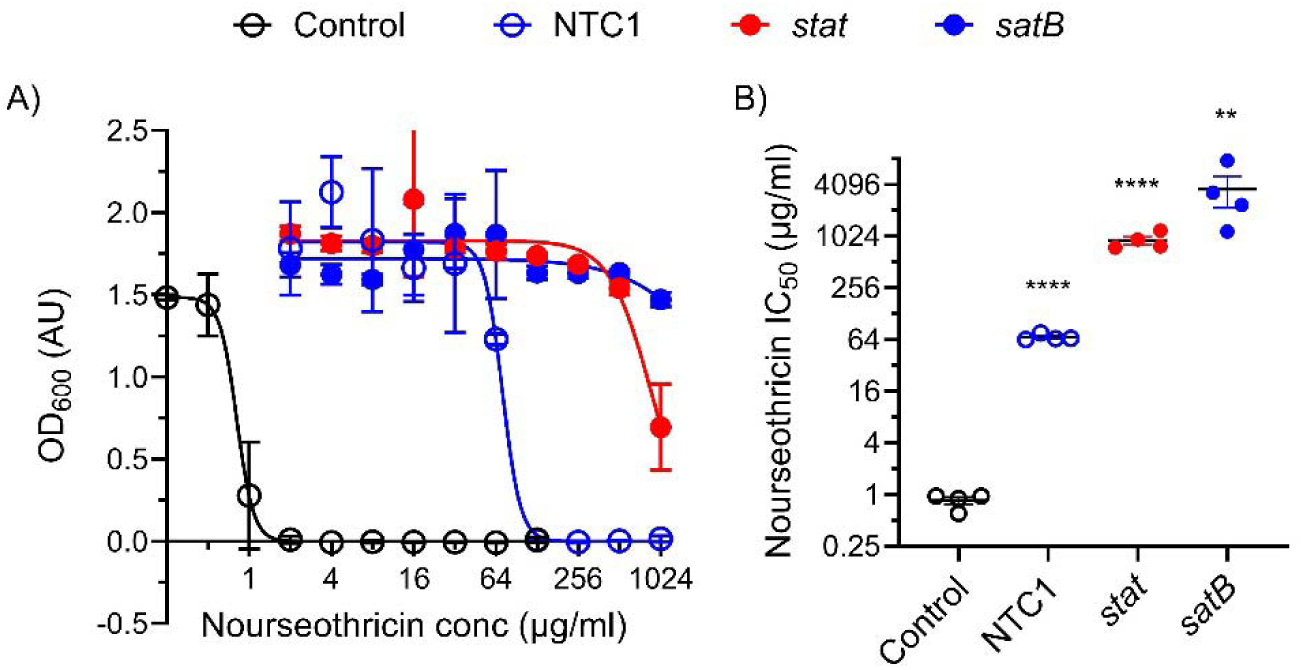
Streptothricin resistance is conferred by the NTC1 metagenomic DNA fragment and *satB*. A) Results of a microbroth dilution assay of *E. coli* clones grown in the presence of variable nourseothricin concentrations (NTC1, *E. coli* carrying the NTC1 metagenomic DNA fragment; *stat*, *E. coli* expressing the *stat* streptothricin acetyltransferase; *satB*, *E. coli* expressing the predicted *satB* open reading frame from the NTC1 fragment). B) IC_50_ values calculated from dose-response curves. (**** p<0.0001, *** p<0.0005, ** p<0.005, n=4 for all).

Given the position of the predicted SatB enzyme on the antimicrobial acetyltransferase phylogenetic tree (**Figure 4B**) and the ability of the *satB* gene to confer streptothricin resistance (**Figure 5**) we next considered our argument that SatB represents a new member of the streptothricin acetyltransferase family. A multiple sequence alignment of SatB and other streptothricin acetyltransferases showed that SatB has identical amino acid residues at predicted key positions for catalysis (Glu137) and streptothricin binding (Leu136, Ala145, Tyr149, Phe154, and Tyr164, numbering based on SatA crystal structure) (**Figure 6A**). A percent amino acid identity matrix of streptothricin acetyltransferases and a single aminoglycoside acetyltransferase indicates that the SatB sequence does not share high levels of identity with any of the other proteins, its highest identity being 37% compared to SAT-4 (**Figure 6B**) (the full-length Needleman-Wunsch identity of SatB with SAT-4 is only 33%). SatB is as unique compared to the other acetyltransferases (average percent identity of 29%) as each acetyltransferase protein is to the rest of the enzymes in the matrix (average percent identity of 29%).

**Figure 6.**
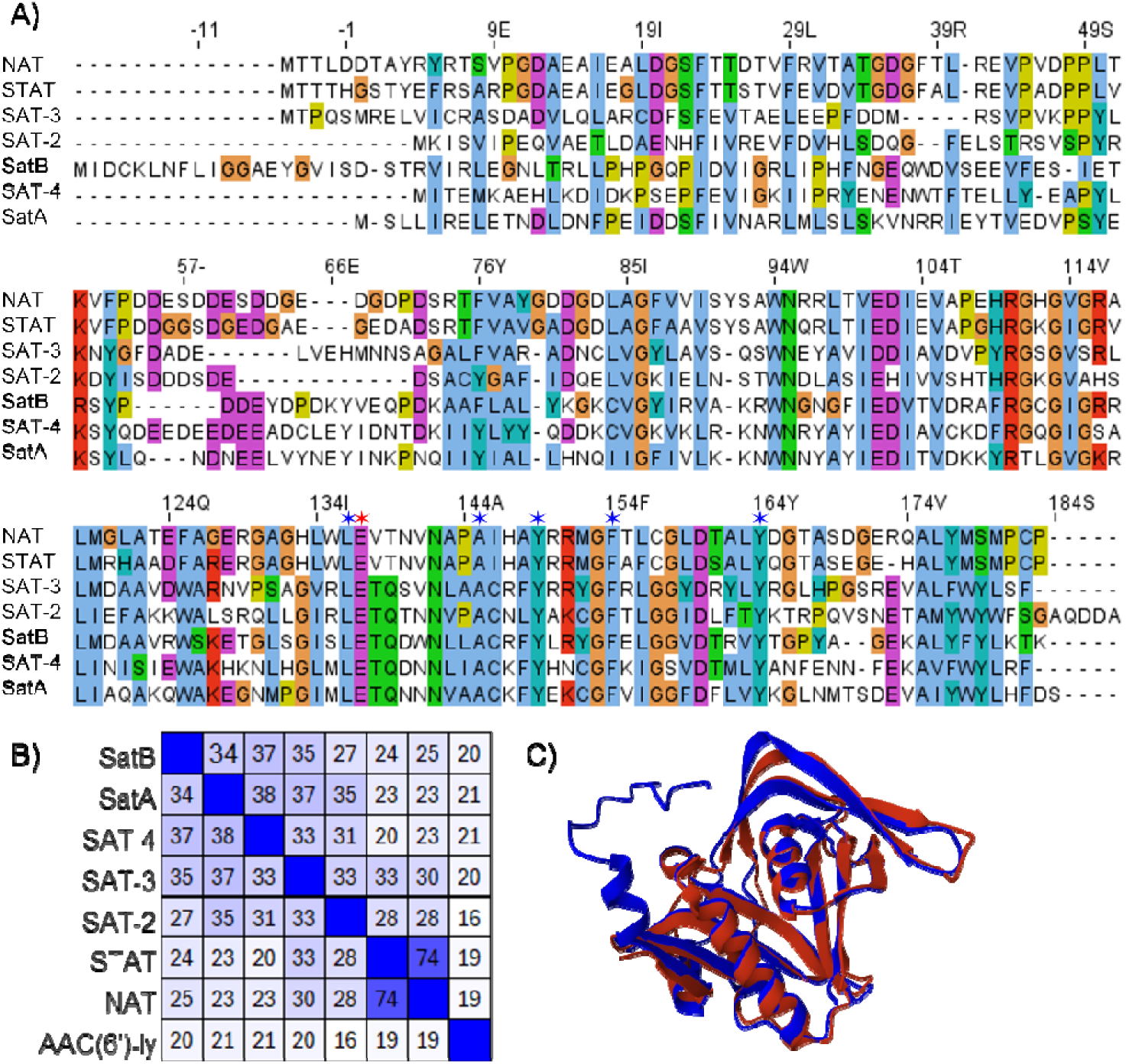
SatB comparison to SatA and other streptothricin acetyltransferases. A) Multiple sequence alignment of the six named streptothricin acetyltransferase enzymes and the proposed SatB enzyme. Conserved residues are highlighted in color, and residues predicted to be important in strepothricin binding are indicated with blue stars (positions 136, 145, 149, 154, and 164) and the predicted catalytic glutamate is highlighted by a red star (position 137) (numbering system corresponds to SatA). B) Heat map of percent amino acid identities between each of the acetyltransferase enzymes, darker colors indicate higher percent identity. C) AlphaFold2 model of the predicted SatB enzyme (blue) superimposed on the SatA crystal structure (red) (PDB 3PP9) (RMSD 2.16 Å).

Finally, we used AlphaFold2^31,32^ to model the three-dimensional structure of SatB and compared it to the solved X-ray crystal structure of SatA from *Bacillus anthracis* (3PP9)^29,33^. When superimposed, the two models show similar secondary structure and have a global root mean squared error (RMSD) of 2.16 Å (**Figure 6C**). In this predicted structure, SatB has an N-terminal region with an alpha helix not found in SatA (**Figure 6C**) which is also evidence in the multiple sequence alignment (**Figure 6A**).

## DISCUSSION

Here we report that the METa assembly method for preparing functional metagenomic libraries can be extended to preparing libraries from low biomass and otherwise precious microbiome samples with limited available metagenomic DNA. As an illustration of this potential, we prepared modestly sized functional metagenomic libraries from metagenomic DNA extracted from an aquarium and from a swab of a human stool sample. We further demonstrated the utility of the resulting functional metagenomic libraries by applying them to identify likely tetracycline efflux pumps and a novel streptothricin acetyltransferase.

A key result of our experiments is that using METa assembly to prepare functional metagenomic libraries allows us move the amplification-free minimum metagenomic DNA input to approximately two orders of magnitude lower than previous protocols^9,15^. An advantage of our tagmentation-based fragmentation approach over physical fragmentation (*e.g., via* sonication) is its ready scalability to lower volumes and therefore lower input DNA masses. However, we suspected that DNA losses during purification and fragment size-selection by gel excision would make sub-200 ng libraries impractical. Instead, we show that, across our experiments, 50 ng or lower input metagenomic DNA masses can be used to prepare functional metagenomic libraries encoding useful amounts of metagenomic DNA (**Table 1**). We propose, and demonstrate, that this opens new sample types to analysis by functional metagenomic selections or screens.

These include samples that are intrinsically of low biomass, either due to difficulty in collection or their rarity (*e.g.,* aquatic metagenomes, but also historically banked microbiomes or samples from remote regions). These also include microbiomes sampled using medical swabs. While our example stool functional metagenomic library used Zymo fecal reference material, the mass of the metagenomic DNA extracted from the swab and used to prepare a functional metagenomic library was less than what has been reported in extractions from other fecal swabs (*i.e.,* ∼100 ng vs ∼370 ng^39^). Functional metagenomic selections have been proposed to be useful for medical applications^42^, and our results support the feasibility of this.

Prior studies that have prepared functional metagenomic libraries from limited metagenomic DNA samples have relied on multiple displacement amplification (MDA) reactions to amplify DNA for input into cosmid large insert functional metagenomic libraries. This approach allowed researchers to convert nanogram or even picogram quantities of metagenomic DNA into the microgram quantities needed for library preparation^43,44^. However, MDA reactions, as well as other amplification-based approaches to increasing DNA mass, come with multiple issues, including selection-based bias (preferential amplification of a subset of templates in the input DNA mixture due to GC content or DNA conformation), drift bias (preferential amplification of a template by random chance), mutation (due to polymerase error present even in high-fidelity enzymes), and risk of amplification of contaminating DNA templates (such as those found in ‘kitomes’)^45,46^. The risk of these biases may be unavoidable for groups preparing functional metagenomic libraries from picogram DNA quantities, but our results show that METa assembly can likely replace MDA for functional metagenomic libraries prepared from nanogram DNA masses.

Our previous smallest METa assembly input, 200 ng, resulted in a functional metagenomic library encoding approximately 13.5 Gb with a preparation efficiency of ∼68 Gb/μg^8^. Here, our 50 ng, 30.5 ng, and 100.8 ng libraries averaged an efficiency of between 20.5 Gb/μg and 24.1 Gb/μg, depending on the inclusion or exclusion of a library that sparked during transformation, suggesting that lower input DNA masses may come with decreased library preparation efficiency (**Table 1**). It is possible that this is due to DNA losses during the multiple purification steps involved in library preparation. Nevertheless, these levels of efficiency still perform more than an order of magnitude better than the average for blunt ligation-based techniques (1 Gb/μg)^8^.

While the functional metagenomic libraries prepared here are smaller than our previous high-mass input libraries, their usefulness is demonstrated by their capture of genes for tetracycline efflux pumps and streptothricin resistance (**Figures 2 and 4**). The three predicted tetracycline efflux pumps show homology to previously identified MFS efflux pumps, with the predicted amino acid sequences having near complete coverage and between 65% and almost 100% identity to hits in the NCBI database. However, they have only low identity to CARD hits, with full length alignments showing only 28% to 29% identity. It is common for proteins identified through functional metagenomic selections for antibiotic resistance to have high identity to proteins in the NCBI database, but for those proteins to not previously have been identified as being antibiotic resistance enzymes^47^. Tetracycline efflux pumps are found within many environments^48–51^ but the difficulty in predicting the substrate specificity of MFS transporters from sequence alone makes the identification of potential resistance genes from environmental microbiomes difficult. This is illustrated by the contrast between the apparent clustering of the TET1 and TET13 MFS pumps with amphenicol efflux and TET3 with mixed macrolide, tetracycline, and other antimicrobial efflux (**Figure 2B**) and their respective lack of biologically relevant resistance to chloramphenicol (**Figure 3CD**) or azithromycin (**Figure 3EF**). Our work highlights the important role that functional metagenomic selections can play in filling this type of knowledge gap.

Our previous work^8,52^ and those of others^41,53^ suggests that streptothricin resistance is common across soil microbiomes and we chose to explore if this is the case for the human gut microbiome as well. The ZymoBIOMICS fecal standard, the source of out stool functional metagenomic library input DNA, has already been sequenced and annotated for known antibiotic resistance genes by the Zymo corporation, but the data associated with this product does not list any known streptothricin resistance genes. Our ability to capture a sequence-novel (**Figure 6**) streptothricin acetyltransferase (termed SatB) from even the modestly sized stool functional metagenomic library suggests that streptothricin resistance may be a feature of animal-associated microbiomes as well. Expression of the *satB* gene in *E. coli* conferred a very high level of nourseothricin resistance (**Figure 5**) and we are planning on testing what effect the extended N-terminal chain of SatB (**Figure 6A and C**) might have on its enzymatic activity and specificity. The presence of this resistance element in the human gut microbiome has important implications for the proposed future use of streptothricin antibiotics in human medicine^54,55^ and suggests that analogs that are not substrates for acetyltransferase enzymes should be explored before clinical adoption of this antimicrobial class^52^.

While we see the opening of amplification-free low biomass functional metagenomic libraries as a significant advancement, limitations to functional metagenomics remain. First, functional metagenomic libraries largely rely on *E. coli* cells as the host organism for the functional metagenomic library and therefore is subject to potential limits on what metagenomic genes can be successfully expressed by this host^56^. In theory, this can be surmounted using alternative hosts^57–59^, and the added efficiency of METa assembly may be important when using strains with lower transformation efficiency than commercial *E. coli* strains. Second, there are potentially diminishing returns to preparing smaller functional metagenomic libraries, although the libraries prepared here were shown to still be useful by their capture of new antibiotic resistance genes even without covering more than a fraction of the genetic diversity found in many microbiomes. Tagmentation reactions are regularly performed on metagenomic DNA samples with 5 ng or less mass^12,60,61^ suggesting that the limit for functional metagenomic libraries could be pushed even further.

In conclusion, while high-throughput DNA sequencing has helped circumvent the great plate count anomaly and has revolutionized microbiome research, methods for accurately annotating novel genes have not managed to keep pace. Machine-learning tools will likely be able to help with this issue but will also require large amounts of high-quality sequence-function correlation data. Functional metagenomics is well placed to supply this data and, in the meantime, can make significant strides in discovering novel functions from known and unknown genes. The fulfillment of this potential by functional metagenomics can be significantly aided by the ability to prepare the libraries more efficiently and using less input DNA, a precondition that we demonstrate is reached here.

## Acknowledgements

CRediT contributions: Conceptualization – TSC; Data curation –TSC; Formal analysis – HMA, EPB, EF, TSC; Funding acquisition – EMH, TSC; Investigation – HMA, EPB, EF, TSC; Project administration – TSC Resources – FJO EMH, TSC; Supervision – EMH, TSC; Visualization – HMA, EPB, EF, TSC; Writing - original draft – HMA, EPB, TSC; Writing - reviewing and editing – HMA, EPB, FJO, EMH, TSC. See supplemental figure 5 for graphical representation of contributions. We would like to thank Chrissy Cabay, William Van Bonn, and James Clark for their help in acquiring the Shedd Aquarium sample and Jinglin Hu for the metagenomic DNA. This work was supported in part by a grant from the Florida State University Council for Research and Creativity.

